# Evaluation of the performance of copy number variant prediction tools for the detection of deletions from whole genome sequencing data

**DOI:** 10.1101/482554

**Authors:** Whitney Whitford, Klaus Lehnert, Russell G. Snell, Jessie C. Jacobsen

## Abstract

**Background:** Whole genome sequencing (WGS) has increased in popularity and decreased in cost over the past decade, rendering this approach as a viable and sensitive method for variant detection. In addition to its utility for single nucleotide variant detection, WGS data has the potential to detect Copy Number Variants (CNV) to fine resolution. Many CNV detection software packages have been developed exploiting four main types of data: read pair, split read, read depth, and assembly based methods. The aim of this study was to evaluate the efficiency of each of these main approaches in detecting deletions.

**Methods:** WGS data and high confidence deletion calls for the individual NA12878 from the Genome in a Bottle consortium were the benchmark dataset. The performance of Breakdancer, CNVnator, Delly, FermiKit, and Pindel was assessed by comparing the accuracy and sensitivity of each software package in detecting deletions exceeding 1kb.

**Results:** There was considerable variability in the outputs of the different WGS CNV detection programs. The best performance was seen from Breakdancer and Delly, with 92.6% and 96.7% sensitivity, respectively and 34.5% and 68.5% false discovery rate (FDR), respectively. In comparison, Pindel, CNVnator, and FermiKit were less effective with sensitivities of 69.1%, 66.0%, and 15.8%, respectively and FDR of 91.3%, 69.0%, and 31.7%, respectively. Concordance across software packages was poor, with only 27 of the total 612 benchmark deletions identified by all five methodologies.

**Conclusions:** The WGS based CNV detection tools evaluated show disparate performance in identifying deletions ≥1kb, particularly those utilising different input data characteristics. Software that exploits read pair based data had the highest sensitivity, namely Breakdancer and Delly. Breakdancer also had the second lowest false discovery rate. Therefore, in this analysis read pair methods (Breakdancer in particular) were the best performing approaches for the identification of deletions ≥1kb, balancing accuracy and sensitivity. There is potential for improvement in the detection algorithms, particularly for reducing FDR. This analysis has validated the utility of WGS based CNV detection software to reliably identify deletions, and these findings will be of use when choosing appropriate software for deletion detection, in both research and diagnostic medicine.

## Introduction

Identifying and characterising genetic variants is central to both genetic medicine and research. Variation is typically categorised into three main classes: single nucleotide variants (SNVs), small insertions and deletions (indels, typically defined as 1-50 bp), and larger structural variants (SVs, typically defined as >1 kb). Structural variants are further subdivided into two categories dependent on whether the change in genetic information is balanced (no gross loss of DNA) or unbalanced (loss or gain of DNA). Deletions and multiplications form the unbalanced copy number variants (CNVs), while translocations or inversions with conservation of the genetic content form balanced chromosomal rearrangements (BCRs). Copy number variants have historically been defined as changes in genetic content >1,000 bp [1]. However, as the resolution of technologies used to identify these variants have improved, this classification has become one of nomenclature rather than practicality, and it is becoming clear that individuals can harbour changes in genetic content in a continuous scale from 1 bp to several Mb.

The study of copy number changes began at a microscopic level in 1959, where visualisation of whole chromosomes (karyotyping) allowed for the first identification of human copy number changes, namely trisomy 21 in Down syndrome patients [2]. This has culminated in attempts to map and catalogue the variation in the human genome represented by copy number variants (CNVs) [3–6] Advances in technology in the intervening decades has resulted in drastic improvement in the ability to detect unbalanced genomic changes, and today partial chromosome changes in genomic content can be detected using multiple techniques. The current diagnostic standard for identifying CNV is chromosomal microarray analysis (CMA). This technology however has limitations in resolution, with clinical reporting thresholds typically >200 kb [7]. Chromosomal microarray analysis reveals the extent of copy number change effecting the region but is unable to resolve the breakpoints to base pair level, and can only detect gross changes in genomic content, not single nucleotide, other small variants, or balanced chromosomal rearrangements.

Decreasing cost has rendered whole genome sequencing (WGS) a viable and sensitive method for CNV detection with rapidly increasing application in genomic research. WGS has the ability of exact base pair resolution; with no theoretical limit on the size of CNV able to be detected. Application of suitable analytical tools has the potential to reveal all types of genetic variation including CNVs, BCRs, indels, and SNVs.

As such, there has been an explosion in the development of software tools to identify CNVs from WGS, with over 80 tools currently available [8]. These tools primarily exploit four different WGS metrics: read depth, split read, read pair, and assembly based, which each rely on distinct information from the sequence data (reviewed in [9, 10]). Briefly, read depth based methods rely upon the theory that the depth of read coverage of a genomic region reflects the relative copy number of the loci, whereby a gain in copy number would result in greater than average coverage. Conversely, a loss in copy number would result in lower than average read coverage of the region. Split read based approaches rely upon paired end sequencing in which only one read from each pair is aligned to the reference genome, while the other one either does not to map or only partially maps to the reference genome. Read pair or paired-end methods exploits discordantly mapped paired-reads where the mapped distance between the read pairs is significantly different from the average fragment size of the library, or if one or both members of the pair is aligned in an unexpected orientation. Finally, unlike the previous approaches which rely on the initial alignment to a reference sequence, assembly based methods *de novo* assemble reads into contigs, which are then aligned and compared to a reference genome.

As each method utilises different information extracted from sequence data, each method has unique strengths and weaknesses. For example, read depth based methods can only identify SVs where there has been an overall change in genetic content (CNVs not BCRs) but confidently detect the direction of genomic change of the CNVs discovered. The performance of read pair methods are reliant on the choice of the alignment algorithm, which can be an issue for repetitive regions due to ambiguities in the correct placement of reads. In general read pair methods are less susceptible to GC bias and are able to identify both CNVs and BCRs. Split read methods require reads that cross the breakpoint and therefore the ability to detect SVs is sensitive to read length. However, this method resolves the breakpoints with single nucleotide accuracy. Finally, the analysis of genome sequence using assembly can have very long run times and require high performance computing resources. This approach does however enable the identification of complex SVs. Taking these points into consideration, a number of software employ a combination of methods to identify CNVs. The performance of WGS-based CNV detection methods has been characterised by fewer reports than those applied to whole exome sequencing based CNV detection [11–15]. However, a recent publication reported the evaluation of read depth-based WGS CNV detection methods and presented a suggested workflow for WGS based CNV detection [16]. Other comparisons of WGS CNV detection methods have been performed in the context of the initial report of the software. Here, we report an unbiased quantitative comparison of the accuracy and sensitivity of all fundamental WGS CNV methodologies for human CNV prediction, testing one representative of each method, and one of the most commonly used combinatorial methods. The resulting performance metrics presented here emphasises the importance of selecting appropriate fit for purpose CNV detection tools.

## Methods

### Sample data

FASTQ and binary alignment map (bam) files aligned to the GRCh37/hg19 reference genome for individual NA12878 was downloaded from the European Nucleotide Archive repository [17].

### WGS-based CNV software

One CNV detection tool was selected from each of the read depth, split read, read pair, and assembly methodologies based on the following criteria: single sample analysis, optimised for high-coverage genomic data (~30-fold coverage), detection of CNVs down to 1kb in size, use in peer-reviewed research, and the software package had to be available to download with a free licence for research/academic use. Based on these criteria, the software packages Breakdancer (v1.4.5) [18], CNVnator (v0.3) [19], Delly (v0.7.7) [20], FermiKit (v0.13) [21], and Pindel (v0.2.5b8) [22] were selected for further analysis (Table 1).

**Table 1.** CNV detection tools used

| Tool | Signals used | SV types detected | Reference |
| --- | --- | --- | --- |
| BreakDancer v1.4.5 | RP | Del, Ins, ITX, Inv, CTX | [18] |
| CNVnator v0.3 | RD | Del, Dup | [19] |
| DELLY v0.7.7 | RP, SR | Del, Ins, ITX, Inv, CTX | [20] |
| FermiKit v0.13 | AS | Del, Ins, Inv, CTX | [21] |
| Pindel v0.2.5b8 | SR | Del, Ins, Inv | [22] |

RP: read pair based, RD: read depth based, SR: split read based, AS: assembly based, Del: deletion, Ins: insertion, ITX: intra-chromosomal translocation, Inv: inversion, CTX: inter-chromosomal translocation.

Each tool was run using the recommended parameters and filtering steps as described in the original publications; CNVnator: 100 for bin size and retaining only variants with a fraction of reads mapped with q0 quality > 0.5, Breakdancer: retaining only variants with a confidence score threshold of Q ≥ 60, Pindel: the number of supporting reads for each CNV was ≥ 2, with Delly and FermiKit using only default parameters with no recommended filtering steps. Comparative evaluation was restricted to deletion calls ≥1kb. More details on the implementation of each tool can be found in **S**upplementary Text 1.

### Deletion detection

The deletion ‘truth-set’ was obtained from the Genome in a Bottle (GIAB) Consortium [23] benchmark SV calls resource, as called by svclassify, a machine learning based approach [24]. This dataset was generated using one-class Support Vector Machines (SVM) where the training data-set was from deletions identified by Personalis Genetics and 1000 Genomes pilot phase deletion calls, and insertions from Spiral Genetics. The 1000 Genomes deletions were called using examples from each methodology: : AB Large Indel Tool, PEMer, BreakDancer, VariationHunter, WTSI, CNVnator, mrFast, Event-Wise-Testing, Pindel, MOSAIK, Cortex, TIGRA, NovelSeq, AbySS, SOAPdenovo, Genome STRiP, and SPANNER. The majority of these calls were independently validated by PCR or array-based experiments. From these deletions, SVM identified annotations that identify CNVs different from random regions of the genome in Illumina HiSeq, PacBio, and Moleculo genome sequence data. The high confidence SVs were therefore called based upon the annotations associated with SVs identified from the machine learning algorithm.

These CNVs have been made available by GIAB to use as a reference standard and have been used in this capacity in a number of studies both in software development and validation [25, 26], as well as the recent read depth WGS deletion detection software evaluation [16]. For this report deletions ≥ 1kb were considered for statistical analysis; consisting of 612 of the 2744 total CNVs reported by GIAB.

The performance of each of the bioinformatic tools was determined by the comparison between the truth-set and predicted deletions generated by the tools. True positives were classified as variants with at least a 50% reciprocal overlap with one or more of the 612 deletions in the filtered GIAB set, as determined by BEDTools (2.26.0)[27]. Concordance between tools was determined as CNVs detected by one or more software, with 50% reciprocal overlap using the python package Intervene [28]

## Results

We selected one software package for each of the four main methodologies of WGS CNV detection and one combinatorial approach for evaluation. Here the predicted deletions from Breakdancer (read pair), CNVnator (read depth), Delly (read pair and split read), FermiKit (assembly), and Pindel (split read) were assessed for accuracy and sensitivity.

### Size distribution of predicted deletions and comparison across tools

We arbitrarily designated bins of eight CNV sizes for investigation of WGS-based variant calls from individual NA12878 (Fig 1). The ‘truth-set’ deletions are relatively evenly divided amongst the bins, with the majority of deletions ≤5 kb (71.6%). Deletions greater than or equal to 1kb predicted by Breakdancer, CNVnator, and FermiKit had a similar size distribution to that of the GIAB identified deletions. In contrast, the outputs from Delly and Pindel were biased towards larger deletions, particularly Pindel with 49.4% >10 kb. This bias is curious given split read based methods are purported to be better suited to detected small deletions than other methodologies [10, 22], however this observation was to extremely small variants (<300 bp).

**Figure 1.**
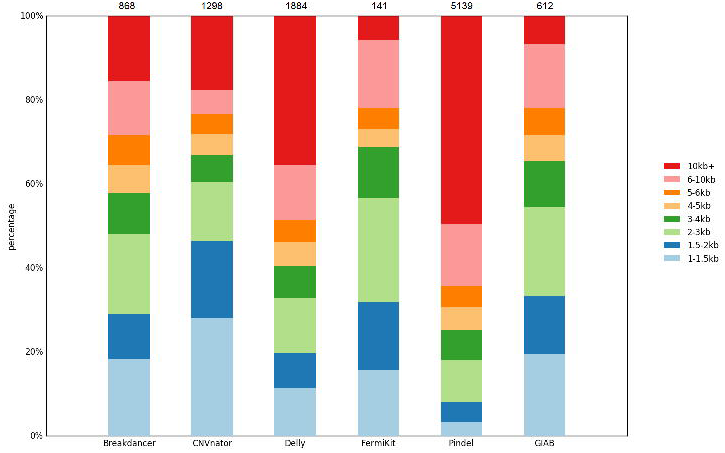
Number and size distribution of deletions ≥1kb predicted by Breakdancer, CNVnator, Delly, FermiKit, and Pindel compared to the GIAB truth-set

### Detection sensitivity of GIAB deletions

The overall performance of each software tool is displayed in Table 2. Viewing the truth-set deletions as a whole, the sensitivity of Delly, Breakdancer, Pindel, CNVnator, and FermiKit, were 96.7%, 92.6%, 69.1%, 66.0%, and 15.8%, respectively. The performance of each tool varied across the size range of deletions (Fig 2A). Pindel displays low sensitivity for small deletions, with 3.29% of deletions correctly identified within the 1-1.5kb range. This is consistent with the low proportion of deletions predicted by Pindel in this size range. The sensitivity for FermiKit was consistently low across all sizes, which likely reflects the small number of deletions predicted by this tool (141 vs 5139 predicted by Pindel, for example). Deletion identification by CNVnator was variable across the size range, without consistent relationship between size and sensitivity. Finally, Breakdancer and Delly performed consistently across the entire size distribution of deletions, with comparable performance between the two tools.

**Table 2.** Deletion detection tool performance

|  | Breakdancer | CNVnator | Delly | FermiKit | Pindel |
| --- | --- | --- | --- | --- | --- |
| Total predicted | 868 | 1300 | 1884 | 141 | 5139 |
| True positives | 567 | 404 | 592 | 97 | 423 |
| False negatives | 45 | 208 | 20 | 515 | 189 |
| False positives | 300 | 898 | 1292 | 45 | 4693 |
| Sensitivity | 0.926 | 0.660 | 0.967 | 0.158 | 0.691 |
| FDR | 0.345 | 0.690 | 0.685 | 0.317 | 0.913 |

Table outlining sensitivity and accuracy of detection for deletions $\geq 1$ kb from individual NA12878 across the five separate bioinformatic tools tested

**Figure 2.**
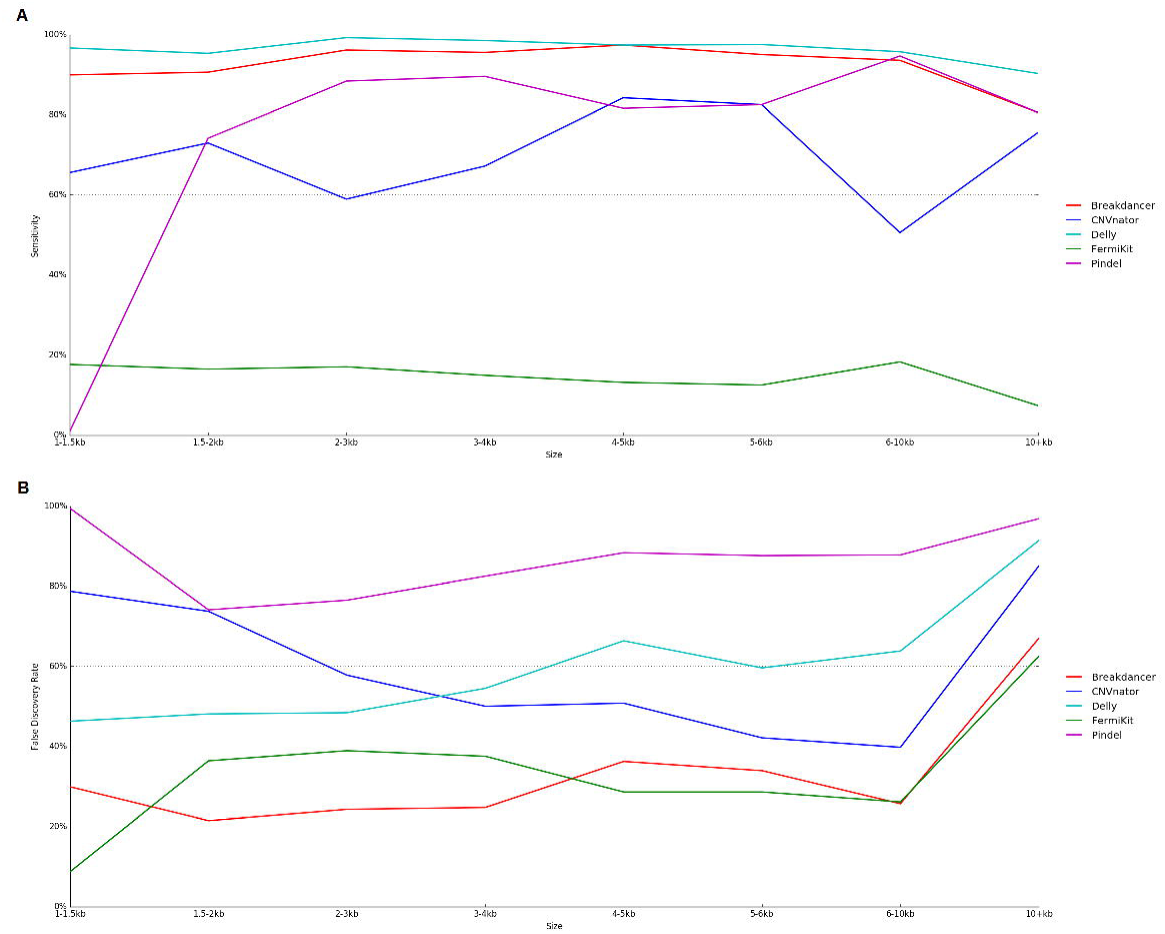
Comparative performance of WGS detection tools across deletion size. (a) Sensitivity of GIAB truth-set (b) False discovery rate of each tool

### Software false discovery rate

There is a natural trade-off between sensitivity and false discovery rate. Often software that delivers a high sensitivity also produce a high proportion of false positive deletions, and thus generate considerable validation work. For deletions ≥1kb, FDR for FermiKit, Breakdancer, Delly, CNVnator, and Pindel were 31.7%, 34.5%, 68.5%, 69.0%, and 91.3%, respectively (Table 2). The FDR for each tool across the deletion size distribution is illustrated in Fig 2B. Pindel had the highest FDR out of all tools across all deletion sizes. The poor performance of Pindel may be due in large part to the substantially higher number of deletions predicted by Pindel than all other tools. Overall, the FDR from CNVnator decreases with increasing deletion size. All tested programs demonstrate FDR > 50% for deletions exceeding 10kb. FDRs for FermiKit and Breakdancer were similar across the deletion size range tested here, while FDR for Delly was consistently higher than that of Fermikit and Breakdancer.

### Concordance of predicted deletions across platforms

There was considerable concordance of correctly identified deletions between individual software packages, where 589 of the 604 correctly identified truth-set deletions where identified by at least 2 tools (Supplementary Table 1). However, the concordance across all packages (Breakdancer, CNVnator, Delly, FermiKit, and Pindel) was relatively poor with only 27 deletions (4.4% of all deletions from the truth-set) identified by all (Fig 3). Only 10 deletions were missed by all tools, the majority of which (60%) were biased towards either end of the size range investigated (>10 kb and 1-1.5 kb in size, Supplementary Figure 1). Relative to the 27 deletions identified by all methods, those missed had fewer interspersed repeat elements (average of 46.59% - including LINES, SINES and Long terminal repeats) compared to 66.57% in the 27 deletions identified by all tools as determined by RepeatMasker.[29].

**Figure 3.**
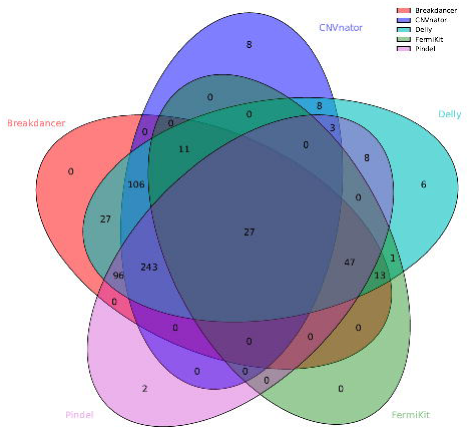
Concordance of deletions ≥1kb correctly identified by WGS software.

The majority (568) of true deletions were identified by Breakdancer and Delly, with very few (36) of the remaining deletions identified by other packages (and not Breakdancer). Of the 568 deletions correctly identified by Breakdancer, all of these were also discovered by Delly (Table S1). In order to investigate if there was extra discriminating power in combining the analysis of Breakdancer and Delly, the number of separately and jointly called false positives was calculated. Only 11 false positive deletions were excluded compared with the application of Breakdancer alone. Thus the FDR for Breakdancer alone (34.5%) showed minor reduction to 33.7% when considering only pairwise deletions with Delly.

## Discussion

The accurate identification of CNVs is important for both research and clinical diagnostics, particularly given CNVs are responsible for the largest percentage of per base genetic variation within genomes (~1.5%) [30]. In comparison, despite the greater number of SNVs per individual (approximately 3.6 million SNVs [31] vs 1,117-1,488 CNVs [30]), they collectively only account for 0.1% of per base genome variation. Chromosomal microarray analysis is the current standard for diagnostic testing for CNVs in human health, however clinical thresholds limit its application to the identification of relatively large scale CNVs of >200kb [7]. Inclusion of WGS for CNV diagnostic testing has the potential to result in a four-fold increase in sensitivity for identification of clinically relevant CNVs (compared to CMA using clinical thresholds alone) [32]. Thus, the increased sensitivity of WGS not only allows for the identification of both small and large scale structural variants, but also enables for the identification of SNV and indels, all within a single test.

A number of CNV detection software packages have been developed which use WGS in various ways including: read depth, split reads, read pair, and assembly based methodologies, or a combination of these methods. However, before WGS CNV detection can be implemented in molecular diagnostics it is necessary to comprehensively assess the methodological performance. A comparative analysis of read depth based methods has recently been reported by Trost, et al., [16]. Here we report an unbiased quantitative comparison of the four primary deletion detection approaches exploiting different features of WGS. We selected Breakdancer (utilising read pair), CNVnator (read depth), Delly (read depth and split read), FermiKit (assembly), and Pindel (split read) software packages for performance assessment. The deletions predicted by each package were compared to high quality deletions (≥1kb) defined in a single individual (NA12878). This dataset has been presented as a reference standard by the Genome in a Bottle (GIAB) Consortium [24], and used in several other studies which quantified the ability of bioinformatic tools to discover CNVs [16, 25, 26]. We found that only Breakdancer and Delly consistently achieved sensitivities over 80% for all deletion sizes, while sensitivities of CNVnator, Pindel, and FermiKit were below 70%. Delly, however had a FDR of 46-91% over the size range of predicted deletions, and the distribution differed to that of GIAB deletions. The deletions predicted by Breakdancer had a distribution that mirrored that of the truth-set deletions, with a relatively low FDR of 21-36% across the size spectrum of deletions.

Concordance analysis of GIAB defined true deletions ≥1 kb for NA12878 discovered between the packages, indicated that there was little overlap between the deletions predicted by the detection software (27 of the total 604 predicted by all packages). Additionally, there was little benefit in combining the best performing tools (Breakdancer and Delly), with only a 0.8% decrease in FDR and a consistent level of sensitivity relative to the performance of the best performing tool (Breakdancer) alone. This was due to no difference in the number of correctly identified deletions when only considering deletions predicted by both packages vs. deletions predicated by Breakdancer alone. Although overall concordance between all software packages was poor, there were a number of deletions predicted by more than one package but not included in the truth-set. Specifically, five deletions were identified by all five packages, but not by the GIAB consortium. Therefore, five loci harbour all characteristics of deletions which can be used to bioinformatically identify deletions from WGS. These loci predicted as deletions by all tools utilising different methodologies whilst not included in the GIAB truth-set indicates the GIAB analyses may not have identified all CNVs in this individual.

The selection of an appropriate truth-set is important for accurate assessment of the performance of biological and bioinformatic tools. Previous comparative CNV studies have utilised results from chromosomal microarray analysis to validate against [12–14], however standard microarrays typically detect CNVs >20 kb making the validation of small CNVs impossible. As many disease-causing deletions reported in the literature are smaller than this threshold, including those identified by this group [33], for this analysis a dataset which included smaller-scale CNVs was required. As such, the GIAB high confidence deletion call set for NA12878 was considered. This is potentially problematic as some of the programs used to generate the calls in the machine learning training dataset are also being evaluated in our analysis (Breakdancer, CNVnator and Pindel). However, these three tools were included in combination with 15 other deletion callers.

To further mitigate confounding impacts, deletions called by Breakdancer, CNVnator and Pindel were only included in the 1000 Genomes dataset if they were confirmed by PCR or array-based experiments. In addition, these deletion calls were only used in the generation of the high-confidence call-set by GIAB to identify the signatures which denote deletions within sequence data from multiple sequencing technologies. Therefore, the potential bias towards identifying deletions called by these software is likely minimal. Interestingly, neither Breakdancer, CNVnator, nor Pindel had the best performance in terms of sensitivity in our analysis, despite their contribution to the development of the training data for the truth-set. Therefore, along with several others [16, 26, 34–36], we deemed this the most appropriate truth set for this analysis, despite some inherent potential biases.

The most accurate method for deletion detection identified in this study is read-pair based as two best performing software packages, Breakdancer and Delly, use this methodology. However, one would assume that the performance of tools incorporating multiple signals would show improved performance in the accuracy and sensitivity of deletion detection. Delly, the only approach tested that used two methods, indeed did show improved sensitivity over the other packages tested. However, Delly also showed a higher FDR across all the size ranges. There is therefore room for improvement in CNV algorithm development, especially in the reduction of the false positive rate, which is particularly important for clinical diagnosis. Determination of the best software package and methodology for the identification of the total scope of SV size (including those less than 1kb) and type (including multiplications and BCRs) will require further investigation.

## Conclusions

WGS based CNV detection tools in this evaluation show widely disparate performance in identifying deletions (≥1kb). Using frequently analysed and comprehensively verified genome alignments for individual NA12878, we conclude that software that exploit read pair-based methods (namely Breakdancer and Delly) showed the highest sensitivity. Of these packages, Breakdancer also had the second lowest false discovery rate over the entire size distribution (34.5%). There was poor concordance in deletions detected by all tools, however there was a large overlap of validated deletions between Breakdancer and Delly. There was little benefit analysing deletions using both packages however, as deletions predicted by both resulted in identical sensitivity (as both packages detected the same number of ‘true’ deletions) and there was only a 0.8% decrease in FDR from 34.5% to 33.7%. While opportunities to improve the detection algorithms remain, primarily reduction of the FDR, read pair-based methods (Breakdancer in particular) are able to effectively identify the majority of deletions and will be of utility as part of bioinformatic pipelines in research and diagnostic medicine.

## Supporting information

## List of abbreviations

BCR: balanced chromosomal rearrangements
CMA: chromosomal microarray analysis
CNV: Copy Number Variant
FDR: false discovery rate
GIAB: Genome in a Bottle consortium
WGS: Whole genome sequencing

## Declarations

### Ethics approval and consent to participate

Not applicable. Humans, animals or plants have not been directly used in this study.

### Consent for publication

Not applicable.

### Availability of data and materials

The genomic sequence data was accessed from the European Nucleotide Archive repository (July 2017), http://www.ebi.ac.uk/ena/data/view/PRJEB1813. Deletion and insertion calls from the Genome in a Bottle (GIAB) Consortium [24] were obtained from ftp://ftp-trace.ncbi.nlm.nih.gov/giab/ftp/technical/svclassify_Manuscript/Supplementary_Information/Personalis_1000_Genomes_deduplicated_deletions.bed and ftp://ftp-trace.ncbi.nlm.nih.gov/giab/ftp/technical/svclassify_Manuscript/Supplementary_Information/Spiral_Genetics_insertions.bed, respectively (July, 2017).

### Competing interests

The authors declare that they have no competing interests.

### Funding

JCJ is supported by a Rutherford Discovery Fellowship from the New Zealand Government and, administered by the Royal Society of New Zealand. The research was funded by the Minds for Minds Charitable Trust and the IHC Foundation.

### Authors’ contributions

WW performed bioinformatic analyses and composed the manuscript. All authors conceptualised the review and experimental methods. JCJ and KL critically reviewed the manuscript. All authors read and approved the final manuscript.

## Acknowledgements

The author(s) wish to acknowledge the contribution of NeSI high-performance computing facilities to the results of this research. New Zealand’s national facilities are provided by the NZ eScience Infrastructure and funded jointly by NeSI’s collaborator institutions and through the Ministry of Business, Innovation & Employment’s Research Infrastructure programme https://www.nesi.org.nz.

---

**Supplementary Figure 1.**
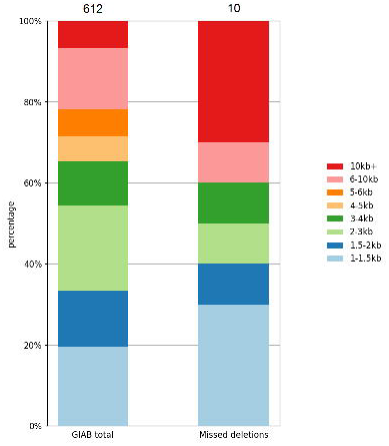
Total number and size distribution of deletions ≥1kb detected in the GIAB truth-set compared to deletions missed by all WGS CNV detection tools in the same truth-set. Total number of deletions in each category listed above.

