## Supplementary material for "Evaluation of the performance of copy number variant prediction tools for the detection of deletions from whole genome sequencing data"

**Supplementary Table 1** Intersect of deletions called WGS based software packages ≥1kb

|  | **Predicted** | **False Positive** | **True Positive** | **False Discovery Rate** |
| --- | --- | --- | --- | --- |
| Pindel | 1245 | 1243 | 2 | 0.998394 |
| FermiKit | 33 | 33 | 0 | 1 |
| FermiKit_Pindel | 0 | 0 | 0 | N/A |
| Delly | 302 | 296 | 6 | 0.980132 |
| Delly_Pindel | 374 | 366 | 8 | 0.97861 |
| Delly_FermiKit | 2 | 1 | 1 | 0.5 |
| Delly_FermiKit_Pindel | 0 | 0 | 0 | N/A |
| CNVnator | 809 | 802 | 7 | 0.991347 |
| CNVnator_Pindel | 5 | 4 | 1 | 0.8 |
| CNVnator_FermiKit | 1 | 1 | 0 | 1 |
| CNVnator_FermiKit_Pindel | 0 | 0 | 0 | N/A |
| CNVnator_Delly | 29 | 21 | 8 | 0.724138 |
| CNVnator_Delly_Pindel | 8 | 5 | 3 | 0.625 |
| CNVnator_Delly_FermiKit | 0 | 0 | 0 | N/A |
| CNVnator_Delly_FermiKit_Pindel | 0 | 0 | 0 | N/A |
| Breakdancer | 10 | 10 | 0 | 1 |
| Breakdancer_Pindel | 1 | 1 | 0 | 1 |
| Breakdancer_FermiKit | 0 | 0 | 0 | N/A |
| Breakdancer_FermiKit_Pindel | 0 | 0 | 0 | N/A |
| Breakdancer_Delly | 98 | 70 | 28 | 0.714286 |
| Breakdancer_Delly_Pindel | 243 | 147 | 96 | 0.604938 |
| Breakdancer_Delly_FermiKit | 13 | 0 | 13 | 0 |
| Breakdancer_Delly_FermiKit_Pindel | 53 | 6 | 47 | 0.113208 |
| Breakdancer_CNVnator | 0 | 0 | 0 | N/A |
| Breakdancer_CNVnator_Pindel | 0 | 0 | 0 | N/A |
| Breakdancer_CNVnator_FermiKit | 0 | 0 | 0 | N/A |
| Breakdancer_CNVnator_FermiKit_Pindel | 0 | 0 | 0 | N/A |
| Breakdancer_CNVnator_Delly | 132 | 27 | 105 | 0.204545 |
| Breakdancer_CNVnator_Delly_Pindel | 276 | 34 | 242 | 0.123188 |
| Breakdancer_CNVnator_Delly_FermiKit | 10 | 0 | 10 | 0 |
| Breakdancer_CNVnator_Delly_FermiKit_Pindel | 32 | 5 | 27 | 0.15625 |

Table outlining the performance of the five tested bioinformatic tools, where as an example FermiKit_Pindel includes variants predicted and identified by these two tools and no others while Breakdancer_CNVnator_Delly_FermiKit_Pindel includes variants predicted and identified by all five tools

**Supplementary Text 1**

**WGS deletion detection description and parameters**

CNVnator utilises the mean-shift approach with multiple-bandwidth partitioning and GC correction in order to predict the presence of CNVs using read depth from aligned WGS data. To generate the initial call set, CNVnator was run concurrently for the whole genome using the bam file as an input, with a bin size of 100. Filtering was then performed, retaining variants with a fraction of reads mapped with q0 quality > 0.5 and which represent deletions ≥1 kb for analysis.

Breakdancer uses the read pair based methodology via the separation distance and alignment orientation between the paired reads versus the insert size distribution estimated from the aligned WGS data. The algorithm identifies SVs from regions with substantially more anomalous read pairs (ARPs) than expected on average. Where the number of ARPs connecting regions exceeds the user-specified threshold, a structural variant is called with a confidence score which is estimated for each variant based on a Poisson model that takes into consideration the number of supporting ARPs, the size of the anchoring regions and the coverage of the genome. The Breakdancer call set was generated for the whole genome using the following default parameters: minimum length of a region [7], cutoff in unit of standard deviation [3], maximum SV size [1000000000], minimum alternative mapping quality [35], minimum number of read pairs required to establish a connection [2], maximum threshold of haploid sequence coverage for regions to be ignored [1000], buffer size for building connection [100], output score filter [30], using the aligned bam as input. From this initial call set, variants were filtered based upon the recommended confidence score threshold of Q ≥ 60 for high coverage WGS, predicted variant type of deletions, and size ≥1 kb.

Pindel uses a pattern growth approach where the software selects the read pairs in which only one of the pairs is mapped to the genome with no mismatched bases from the alignment. Pindel then determines the anchor point and orientation of the unpaired split-read. In this study, Pindel was run on the whole genome concurrently with a pindel file generated using the included bam2pindel.pl Perl script from the NA12878 aligned bam. The following parameters were used for variant calling: window_size [10 million], max range index [5], report close mapped reads [false], min NT size [50], min inversion size [50 bp], min num matched bases [30], additional (site) mismatch [1], min perfect match around BP [3], sequencing error rate the expected fraction of sequencing errors [0.05], maximum allowed mismatch rate [0.1]. From the output, these calls were filtered to retain variants in which the number of supporting reads for each CNV was ≥ 2, that were called as deletions, and those with a predicted size ≥1 kb.

Delly analysis consists of two separate components, separately utilising both paired-end and split-read data. Paired-end analysis involves the identification of read pairs which are outliers on the insert size distribution or the pairs that have an unexpected orientation. Delly does not have adjustable parameters for user input. Therefore for this investigation, deletions were called separately on a genome wide scale using Delly from the aligned bam for NA12878. From those called by Delly, deletions were filtered based on size ≥1 kb.

FermiKit can be classified as an assembly based method for SV detection. This software performs a *de novo* assembly of the reads and then maps this assembly to a reference genome to call variants. FermiKit inherently lacks user defined parameters outside of informing the algorithm of the size of the genome and read length. For this study variants called by FermiKit were run with genome size [3g] and read length [150] from the NA12878 fastq input. Similar to all other CNV calling programs, from the variants called by FermiKit, filtering was performed to retain deletions ≥1 kb.

CNVnator, Pindel and Delly also estimate zygosity of the predicted CNVs either natively, or through the included functionality to covert the output to VCF. However as not all of the tools analysed here incorporate zygosity calls, this was excluded from the analysis.
